# Macronutrient composition of Button Grass, Curly Mitchell Grass, Native Millet and Weeping Grass grains for their potential in modern food applications

**DOI:** 10.1101/2025.10.13.682222

**Authors:** Farkhondeh Abedi, Claudia Keitel, Ali Khoddami, Angela L. Pattison, Thomas H. Roberts

## Abstract

Grains of Australian native grasses have been important components of traditional Aboriginal diets for millennia and have the potential for increased utilisation in contemporary food systems. This study assessed the macronutrient profiles of whole grains from *Dactyloctenium radulans* (Button Grass), *Astrebla lappacea* (Curly Mitchell Grass), *Panicum decompositum* (Native Millet), and *Microlaena stipoides* (Weeping Grass) compared to wheat, barley, and sorghum using proximate analysis, Osborne protein fractionation, gel electrophoresis, and carbohydrate assays. Key results were that Native Millet had high lipid content (8.0 g/100 g dry weight basis (db)), Curly Mitchell Grass had high protein (29.1 g/100 g db) and low carbohydrate content (64.0 g/100 g db), and there were substantial differences in prolamin and glutelin fractions across the species. All four native grains were gluten-free, and their starch amylose content ranged from 25.7% (Button Grass) to 41.2% (Curly Mitchell Grass), which affects starch properties. Button Grass had the highest dietary fibre content (21.9 g/100 g db), while Weeping Grass had the highest beta-glucan levels (8.6 g/100 g db), supporting functional food applications. Our findings highlight the potential for an expanded range of food applications for these grains and their contribution to human nutrition, together with implications for supporting Indigenous-led enterprises.

## 1. Introduction

Over 1,100 Australian native grass species have been recognized (Drake et al., 2021), many of which have deep root systems and are tolerant to high temperatures and drought. They have integrated into Australia’s natural biodiversity over millions of years of evolution. Seeds from certain grasses have held significance in the traditional diets of First Nations people in Australia for tens of thousands of years, varying with specific regions and Indigenous cultures (Pascoe, 2014).

The global cereal grain industry faces the complex task of providing food for a growing population while minimizing environmental impact. Australian native grains, often overlooked in agricultural settings, present an opportunity to help address these challenges (Sydney Institute of Agriculture, 2020). As consumers increasingly seek sustainably acquired and nutritious products, the integration of Australian native grains into the industry can meet this demand, promoting a more diverse and adaptable food supply (Andreotti et al., 2022; Fernández-Tomé et al., 2023; Jenifer et al., 2023; Kaur et al., 2025).

Together, the cultivation and processing (including threshing) of native grains for food applications have the capacity to revitalise cultural practices, create new economic opportunities, promote local agriculture, and foster entrepreneurship within Indigenous communities (Jenifer et al., 2023). To assist Indigenous-led enterprises to create native grain-based foods that align with consumer preferences for healthier, diverse, and sustainably sourced products, a thorough understanding of the macronutrient composition of these grains is required. Quantifying the macronutrient profiles of Australian native grains helps to position these ingredients in various food applications as either superior candidates compared to their domesticated counterparts, or partial substitutes for domesticated grains.

However, their broader adoption is constrained by yield variability and generally lower productivity. Native species yield between 8 and 2,200 kg per hectare, depending on species and conditions, whereas commercial wheat can produce up to 10 tonnes ha^-1^ in high-rainfall areas (Cole & Johnston, 2006; Drake et al., 2021). This difference in yield presents a major challenge for the integration of native grains into conventional agri-food systems.

In 2023, we completed a study of the macrostructure, microstructure and histochemistry of the grains of four Australian native grass species: *Dactyloctenium radulans* (Button Grass)*, Astrebla lappacea* (Curly Mitchell Grass)*, Panicum decompositum* (Native Millet), and *Microlaena stipoides* (Weeping Grass) (Abedi et al., 2023). These species were selected based on their importance in First Nations people’s traditional diets, their capacity for efficient production and processing, and their potential for modern applications in the food industry. The results were compared to those obtained for three domesticated control grains: red sorghum (*Sorghum bicolor*), 2-row spring barley (*Hordeum vulgare*) and bread wheat (*Triticum aestivum*). Substantial differences in the size, shape and colour of the grains, the size of the embryo as a proportion of the whole grain, and the thickness of the aleurone layer were found among the four native grasses.

There is a limited number of publications available on the biochemical properties of Australian native grains (Birch et al., 2023; Brand-Miller & Holt, 1998; Cowley et al., 2023; Foster et al., 2010; Williams et al., 2024). Building on these papers and on our grain structure and histochemistry study discussed above, the overall objective of this study was to determine the macronutrient composition of four Australian native grains. We performed biochemical analyses of grains of the same species as those studied in Abedi et al. (2023). These experiments comprised a proximate analysis, Osborne fractionation and sodium dodecyl sulfate-polyacrylamide gel electrophoresis (SDS-PAGE) of the proteins, a sensitive gluten test, as well as carbohydrate analyses, namely total starch, amylose, total dietary fibre, beta-glucan, and resistant starch assays.

This study was conducted in partnership with the Gamilaraay Grains Custodians (GGC), who are members of the Gamilaraay/Yuwalaraay Indigenous communities in northern NSW. The GGC provide cultural authority to help direct the vision and strategy of the revitalisation of Gamilaraay grains with the option to co-design and co-produce aims, methods, outcomes and communications for native grains research and education at the University of Sydney’s Plant Breeding Institute at Narrabri. The GGC guide native grains knowledge and activities that benefit local communities. Consultation with the GGC regarding the current manuscript was conducted via a Zoom meeting on 12 August 2024. A copy of the mature manuscript was then made available to the GGC for comment via a protected website from 13 August 2024.

## 2. Materials and methods

Most analyses were performed in triplicate using whole-grain flour prepared from the same batch of grains for each species, with each batch derived from multiple plants. Moisture and total lipid measurements were conducted in duplicate due to limited sample availability.

All values are expressed on a dry weight basis (db), except when otherwise indicated (i.e. on a wet weight basis (wb) to allow for comparison with literature values for which moisture contents were not determined.

### 2.1. Plant materials

We examined grains of Button Grass (*Dactyloctenium radulans*), Curly Mitchell Grass (*Astrebla lappacea*), Native Millet (*Panicum decompositum*), and Weeping Grass (*Microlaena stipoides*). Grains of sorghum (*Sorghum bicolor* cv. Buster) grown at Narrabri, New South Wales (NSW), barley (*Hordeum vulgare* cv. Spartacus) grown at Hanwood, NSW, and bread wheat (*Triticum aestivum* cv. Kord) grown at Wasleys, South Australia were employed as control samples.

Button Grass and Native Millet seeds were hand-harvested from the University of Sydney Plant Breeding Institute, and Llara farm, respectively, both situated near Narrabri (latitude: -30° 18’ S; longitude: 149° 45’ E) in northern NSW, Australia.

Curly Mitchell Grass seeds were harvested on a farm near Thallon (latitude: -28° 38’ S; longitude: 148° 51’ E), Queensland, while Creative Native Food Service Co, Australia, provided the Weeping Grass seeds, which were harvested from a farm outside Armidale (latitude: -30° 30’ S; longitude: 151° 40’ E), NSW.

Of the four native grains studied here, only the Native Millet samples included the palea and lemma. Otherwise, ‘grain’ is defined botanically as the caryopsis, an indehiscent fruit commonly observed in grasses, which is distinguished by its enclosed structure at maturity. The caryopsis comprises the true seed along with its testa and pericarp, with the absence of palea and lemma components.

All the grains sourced for this study were obtained in May 2021. Threshing methods suitable for each species were followed to prepare the grains for analysis, as outlined in Pattison et al. (2023). The cleaned grains were then stored at 4°C until further analysis.

### 2.2. Flour preparation

Whole-grain samples were ground using a Laboratory Mill 3100 (Perten, Australia) with a 0.5-mm screen size before analysis.

### 2.3. Proximate analysis

Proximate analysis of the native and control grains was conducted to determine their moisture, protein, lipid, ash and carbohydrate content (the latter by difference).

#### 2.3.1. Moisture

The moisture content in the native and control grains was measured following the AACC 44–15.02 method (American Association of Cereal Chemists-International, 2000). For this process, 2-g samples were placed into previously weighed aluminium dishes. After being covered and accurately weighed, the dishes were placed without their covers, alongside the lids, into an oven at 135 ± 1°C for 2 h. The timing began once the oven had reheated back to 135°C, which took ∼2 min. The dishes were then removed from the oven, immediately covered, and set in a desiccator to cool to room temperature (for ∼1 h), before their weight was measured. The following formula was used to calculate the moisture content: Moisture (%) = (A / B) ×100, where A represents the loss of moisture in grams, and B is the initial weight of the sample in grams.

#### 2.3.2. Total carbohydrate

The percentages of moisture, ash, lipid, and protein were summed and subtracted from 100% to determine total carbohydrate content.

#### 2.3.3. Total protein

The total protein content of the grain samples was measured using the Dumas combustion method AACC 46-30.01. An Elemental Analyser (Vario CHN Cube, Elementar, Germany) was employed to determine the nitrogen content of the samples.

For the calculation of total protein content, a conversion factor of 5.8 was employed for both the Australian native grains and control samples. While 6.25 is the general factor traditionally applied across food matrices, it assumes that the proteins in the food contain 16% nitrogen, which does not accurately reflect the amino acid composition of cereal proteins. The factor 5.8 is more appropriate for proteins of cereal grains such as wheat and barley, which typically contain lower nitrogen percentages (Mariotti et al., 2008). Although species-specific factors for the native grains are not yet available, using 5.8 across all samples ensures consistency and aligns with accepted practices in cereal grain protein analysis.

#### 2.3.4. Total lipid

The total lipid content of the native and control grain species was measured using Soxhlet extraction, following the principles outlined in the AOAC 920.39C Method for Cereal Fat (Association of Official Analytical Chemists). Approximately 2 g of each grain flour sample was accurately weighed and placed into a Soxhlet thimble. Hexane was added to a pre-weighed, dry round-bottom flask until the solvent level allowed for two siphon cycles. The heated solvent surrounded the sample for 5–10 min before siphoning back into the boiling flask. This cycle was maintained for 4 h, after which the hexane containing the extracted fat was allowed to evaporate to ensure complete solvent removal. The round-bottom flask, now containing only the extracted fat, was re-weighed. The total fat content of each sample was calculated using the difference in the weight of the pre-weighed flask before and after extraction.

#### 2.3.5. Ash

Ash content was measured as part of the dietary fibre analysis (see 2.5.3). After extracting dietary fibre, the residue was isolated and placed in crucibles for ash content determination. The ash content, which was subtracted from the total residue weight for the dietary fibre calculation, was quantified by incinerating the samples. The crucibles containing the residue were placed in a muffle furnace at 550°C for 5 h. After incineration, the crucibles were cooled to room temperature in a desiccator and then re-weighed. The ash content was calculated by subtracting the initial weight of the crucible and sample from their combined weight after incineration.

### 2.4. Protein analysis

#### 2.4.1. Protein fractionation via the Osborne method

A modified Osborne fractionation procedure was conducted to extract proteins from the native grains and barley (as control) whole-grain flours, based on Osborne’s 1907 method (Osborne, 1907). In each extraction, 1 g of the flour was suspended in 10 mL of Milli-Q water to solubilise the albumins and mixed for 30 min at 100 revolutions per minute (rpm) using an Intelli-mixer RM-2 (John Morris, Australia) in “UU” mode, followed by centrifugation at 2,600 *g* for 20 min using an Eppendorf centrifuge 5810/5810 R. The pellet was then dissolved in another 10 mL of Milli-Q water and mixed and centrifuged as above.

The steps above were repeated a third and fourth time. Following the semi-exhaustive extraction of the albumins, the pellet was suspended in 10 mL of 0.4 mol L ¹ sodium chloride solution to solubilise the globulins, and the mixing and centrifugation steps were repeated three more times. After the final round of salt extraction, the supernatant was retained, and the pellet suspended in 10 mL of 70% (v/v) ethanol to solubilise the prolamins, with the same mixing and centrifugation procedure applied.

Finally, the procedure was repeated with 10 mL of 0.1 mol L ¹ sodium hydroxide added to the pellet to solubilise the glutelins. In each of the four rounds of extraction for the four types of solvent, the resulting supernatants were retained and then pooled for protein content determination using the bicinchoninic acid method (BCA assay, Thermo Scientific Pierce^TM^, Rockford, IL, USA) following the manufacturer’s instructions, employing a UV-1600PC spectrophotometer. Thus, estimates of the absolute and relative abundances of albumins, globulins, prolamins, and glutelins in the whole-grain flour samples were obtained (Van de Vondel et al., 2020).

#### 2.4.2. Characterisation of the protein profiles of each Osborne fraction

Protein profiles of each Osborne fraction were analysed using SDS-PAGE. Albumin, globulin, prolamin and glutelin fractions extracted from the native grains and barley were diluted to the lowest protein fraction concentration (the globulin concentration in Button Grass: 0.263 mg mL^-1^). SDS-PAGE was performed using a BioRad Mini-PROTEAN Tetra Vertical Electrophoresis Cell apparatus and 4–20% precast polyacrylamide Mini-PROTEAN TGX gels (1 mm thick; Bio-Rad Laboratories, USA). The gels were run at 200 V (constant voltage) for 35 min until the bromophenol blue marker had reached the bottom of the gels. Replicate gels were stained with either Coomassie Brilliant Blue R-250 or silver nitrate.

For the Coomassie Brilliant Blue staining, the staining and destaining protocol used involved two main solutions: the Super Destaining Solution and Coomassie Brilliant Blue R-250 dye. The Super Destaining Solution was composed of glacial acetic acid, methanol, and Milli-Q water in a ratio of 1:4:5 (v/v/v). To prepare 100 mL of Coomassie Brilliant Blue R-250 dye, 100 mg of Coomassie R-250 dye was added to 90 mL of the Super Destaining Solution and mixed thoroughly with a magnetic stirrer until the dye was completely dissolved.

The gel was immersed in the Brilliant Blue R-250 Staining Solution and then subjected to microwave heating at high power until boiling (30–60 s). Subsequently, the stained gel was destained by microwave heating at high power (30–60 s) in the Super Destaining Solution until boiling. The gel was further incubated in the Super Destaining Solution for 1 h to enhance band visualisation.

For the silver nitrate staining, the gels were incubated in a solution of 25% ethanol, 1% HNO_3_, and 0.2% AgNO_3_ for 5–10 min with gentle agitation. The gels were then rinsed with distilled water for 3 min to remove excess stain and impurities and incubated in the development solution (3% Na_2_CO_3_ and 0.2% HCOH) for 2–5 min at room temperature with gentle agitation. In the last step, the gels were incubated in distilled water for 2 min to halt the staining process.

Images of the gels were recorded using a ChemiDoc MP imaging system. Background subtraction for the gel images was conducted using ImageJ software. Additionally, brightness and contrast adjustments were applied using the “Auto” function in ImageJ to enhance visualisation.

#### 2.4.3. Gluten screening

Native and control grains (including brown rice) were analysed using a 3M™ Gluten Protein Rapid Kit according to the manufacturer’s instructions. This kit employs a lateral flow device (LFD) based on an immunochromatographic test method utilising a polyclonal antibody specifically designed for detecting gliadin in wheat, rye (secalins), and barley (hordeins) proteins. The ground sample (0.2 g) was transferred to a microcentrifuge tube. Extraction Buffer (1.8 mL of 3 mol L ¹) was added, followed by vigorous vortexing for 3 min.

The extracted sample was centrifuged for 20–30 s at 2,350–4,600 *g*, with the supernatant collected as the extracted sample. An aliquot (100 μL) of the extracted sample was transferred to the designated sample well on the LFD (the wheat sample was diluted in a 1:3 ratio with the extraction buffer due to its high viscosity). After 10 min, the sample was regarded as positive for gluten if both the test and the control lines were visible on the LFD. If only the control line was visible on the LFD, the sample was considered gluten-free. Additionally, any faint appearance of the test line was interpreted as a negative result, indicating the absence of detectable levels of gluten protein in the sample. The images were recorded employing a Nikon D3400 digital single-lens reflex (DSLR) camera (designed by Nikon Corporation in Japan, manufactured in Thailand).

### 2.5. Carbohydrate characterisation

#### 2.5.1. Total starch

For total starch content determination, a Total Starch Assay Kit (AA/AMG) from Megazyme (Wicklow, Ireland) was employed. The method involves hydrolytic digestion with α-amylase (AA) and amyloglucosidase (AMG), converting starch to maltodextrins and D-glucose, respectively. This is followed by the use of glucose oxidase-peroxidase (GOPOD) reagent and the measurement of absorbance at 510 nm to quantify the D-glucose using a Shimadzu UV 1900 spectrophotometer (McCleary et al., 1997).

#### 2.5.2. Amylose content

The percentage of amylose in the total starch content of the samples was determined using an Amylose/Amylopectin Megazyme Assay Kit from Megazyme. The method employed lectin concanavalin A (Con A) to precipitate and remove the amylopectin-Con A complex. Subsequently, the remaining amylose molecules were hydrolysed to D-glucose, using AA/AMG, after which D-glucose was measured spectrophotometrically at 510 nm with GOPOD reagent using a Shimadzu UV 1900 spectrophotometer (Yun & Matheson, 1990).

#### 2.5.3. Total dietary fibre

The dietary fibre content of the whole-grain flours was determined through the enzymatic/gravimetric method outlined by Prosky et al. (1988), employing a Total Dietary Fiber Megazyme Assay Kit from Megazyme. AA was used to induce gelatinisation, hydrolysis, and depolymerisation of the starch. Protease and AMG were then added to the sample. Soluble fibre was precipitated, and depolymerised proteins and glucose derived from the starch were eliminated. The residue was filtered, dried, and weighed. The tests were run in duplicate for each sample; one of the duplicate samples was subjected to protein analysis, the other for ash determination. The total dietary fibre was calculated by subtracting the combined weight of the protein and ash from the weight of the filtered and dried residue.

#### 2.5.4. Resistant starch content

The resistant starch (RS) content of the flours was determined using a Resistant Starch Rapid Kit from Megazyme. Pancreatic AA and AMG were employed to solubilise and hydrolyse non-resistant starch into D-glucose. RS was recovered as a pellet by centrifugation. The pellet was dissolved in 1.7 mol L ¹ NaOH and neutralised with acetate buffer (pH 3.8). AMG was utilised to hydrolyse the starch to D-glucose, followed by the use of GOPOD reagent and reading the absorbance at 510 nm to determine the D-glucose using a Shimadzu UV 1900 spectrophotometer (McCleary et al., 2002).

#### 2.5.6. Beta-glucan content

For beta-glucan (β-D-glucan) measurement, a Mixed Linkage β-glucan Kit from Megazyme was used. Briefly, samples were incubated with lichenase enzyme and filtered, followed by hydrolysis with β-D-glucosidase. Lichenase specifically breaks down β-D-glucan into oligosaccharides, which are further hydrolysed into glucose by β-glucosidase. The sample was reacted with GOPOD reagent, and the absorbance was measured at 510 nm using a Shimadzu UV 1900 spectrophotometer to determine glucose concentration. (McCleary & Codd, 1991).

### 2.6. Statistical analysis

For sample characterisation, most measurements were performed in triplicate for each flour sample from each of the species. This denotes that three separate measurements were taken for each individual flour sample, and the process was repeated for all species in the study. Moisture and total lipid measurements, however, were performed in duplicate, and the mean of the two values was used for statistical analysis. The standard deviation was calculated accordingly to reflect variability between the replicates. All data represent the mean ± standard deviation. Data were analysed using GraphPad Prism v.10.2.1 (GraphPad Software, San Diego, CA, USA). Standard one-way analysis of variance (ANOVA) was conducted to assess differences among the tested groups for all measured parameters. Tukey’s *post hoc* multiple comparisons test was performed to identify significant differences between groups. Statistical significance was determined at p ≤ 0.05.

## 3. Results and Discussion

### 3.1. Proximate analysis

#### 3.1.1. Moisture

Moisture content, which was determined to allow the other grain components to be expressed on a dry weight basis, was in the range of 10.2–14.7% (Table 1). These values compare well with Australian moisture content receival standards; e.g., 12.5% for milling wheat and malting barley; 13.5% for Australian White Wheat (AWW) and feed barley (CBH Group ‘Moisture Management’ website).

**Table 1.** Proximate analysis of the studied native and control grains. The moisture content was reported as a percentage of the total grain weight. Total carbohydrate, protein, lipid, and ash contents were reported on a dry weight basis (g/100 g). The total carbohydrate content was calculated by difference.

| Species | Moisture | Total carbohydrate | Total protein | Total lipid | Ash |
| --- | --- | --- | --- | --- | --- |
| Button Grass | 12.3 ± 0.4<sup>b, c</sup> | 74.1 ± 0.4 | 14.4 ± 0.3<sup>c, d</sup> | 2.8 ± 0.1<sup>d</sup> | 8.7 ± 0.3<sup>a</sup> |
| Curly Mitchell Grass | 10.2 ± 0.6<sup>d</sup> | 64.0 ± 0.5 | 29.1 ± 0.2<sup>a</sup> | 3.6 ± 0.4<sup>c</sup> | 3.3 ± 0.2<sup>b</sup> |
| Native Millet | 14.7 ± 0.6<sup>a</sup> | 71.9 ± 1.0 | 16.7 ± 0.7<sup>b</sup> | 8.0 ± 0.6<sup>a</sup> | 3.4 ± 0.4<sup>b</sup> |
| Weeping Grass | 13.5 ± 0.7<sup>a, b</sup> | 80.0 ± 0.4 | 15.5 ± 0.2<sup>b, c</sup> | 2.0 ± 0.4<sup>e</sup> | 2.5 ± 0.1<sup>c</sup> |
| Wheat | 11.3 ± 1.2<sup>c, d</sup> | 82.7 ± 0.4 | 12.5 ± 0.1<sup>e</sup> | 2.5 ± 0.4<sup>d, e</sup> | 2.2 ± 0.2<sup>c</sup> |
| Barley | 13.3 ± 0.6<sup>b</sup> | 81.7 ± 0.2 | 10.7 ± 0.1<sup>f</sup> | 4.1 ± 0.1<sup>c</sup> | 3.5 ± 0.2<sup>b</sup> |
| Sorghum | 11.0 ± 0.1<sup>c, d</sup> | 78.9 ± 1.2 | 13.2 ± 1.1<sup>d, e</sup> | 5.0 ± 0.0<sup>b</sup> | 2.9 ± 0.3<sup>b, c</sup> |

Data are presented as mean ± standard deviation. Total protein and ash determinations were conducted in triplicate (n=3), while total lipid and moisture determinations were conducted in duplicate (n=2), using wholemeal flour from the same batch of grains. Means with different letters are significantly different ( $p \leq 0.05$ ), as determined by Tukey's multiple comparisons test following one-way ANOVA.

In comparison, teff (*Eragrostis tef*) was reported to contain 8.82% moisture, which is lower than the moisture range observed in the native grains studied here (Zhu, 2018). Similarly, browntop millet was reported to have a moisture content of 8.69%, also below the native grain values (Ponnapalli et al., 2023). However, variations in grain type, harvest maturity, and post-harvest handling may influence these differences.

#### 3.1.2. Total carbohydrate

Total carbohydrate content was the highest in Weeping Grass (80.0 ± 0.4 g/100 g) and Button Grass (74.1 ± 4.0 g/100 g) among the native grains (Table 1). Curly Mitchell Grass had the lowest carbohydrate content at 64.0 ± 0.5 g/100 g among both the native and control grains (Table 1). Considering its low total carbohydrate content compared to wheat and barley (Table 1), Curly Mitchell Grass might be particularly suitable for low-carbohydrate baked goods, diabetic-friendly foods, and weight management products (Gasparre et al., 2024).

The total carbohydrate content of teff (*Eragrostis tef*) was reported as 73.13 g/100 g (Zhu, 2018), placing it close to Button Grass and slightly lower than Weeping Grass, but considerably higher than Curly Mitchell Grass.

#### 3.1.3. Total protein

Total protein contents across the studied native grain species (Table 1; Fig. 1) were consistent with those of studies by Brand-Miller et al. (1998), Foster et al. (2010) and Birch et al. (2023). Grains of Curly Mitchell Grass had a protein content roughly double that of the other native species and triple that of barley (*p* ≤ 0.001). This result is consistent with the findings of Foster et al. (2010), who reported a value of 29.1 g/100 g for the same species. It is also comparable to the 24.68 g/100 g reported by Williams et al. (2024) for the total protein content of an unspecified Australian native grain.

**Figure 1.**
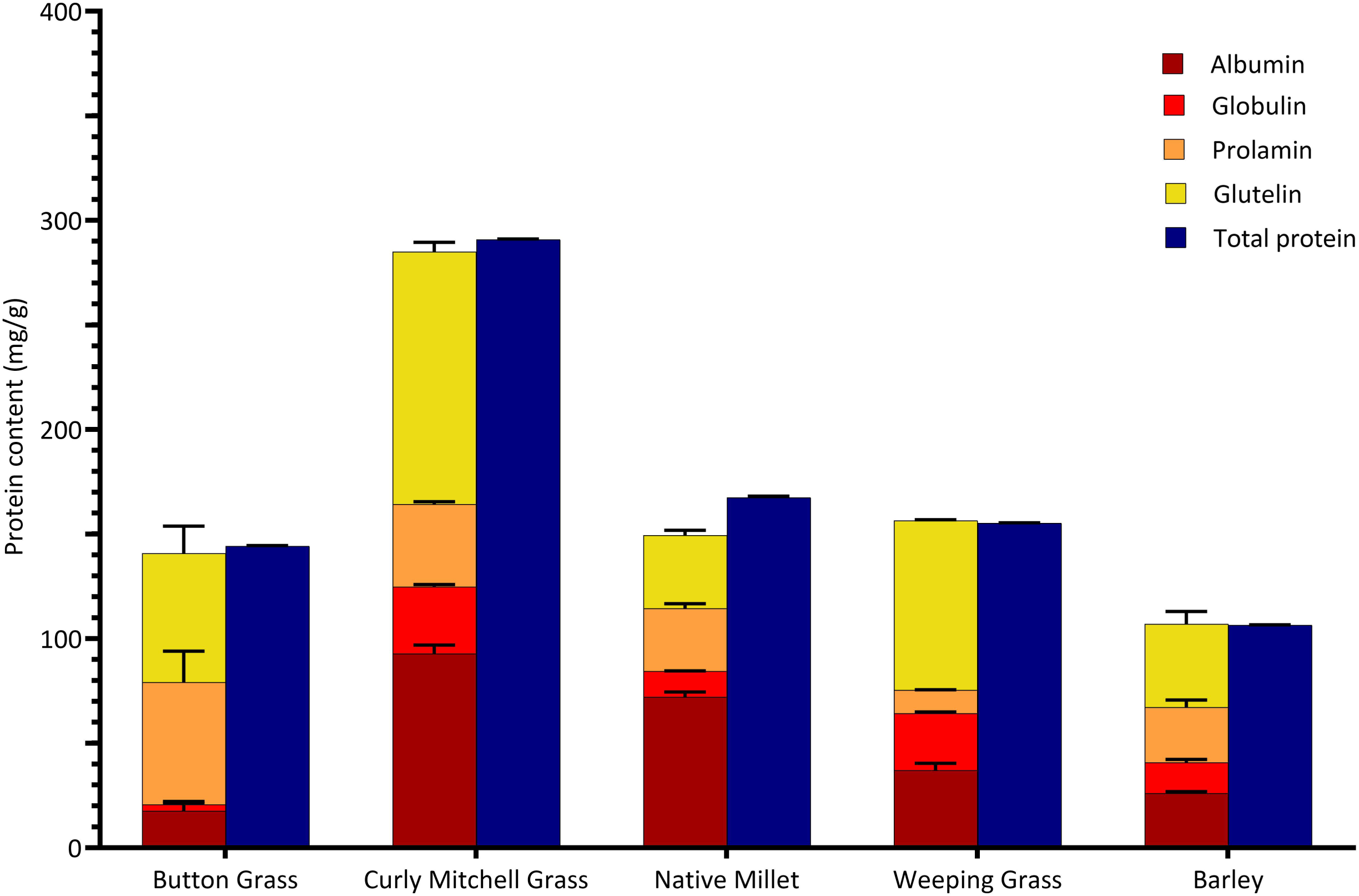
Mean concentrations of the protein fractions in the native grains and barley. Albumin (soluble in water), globulin (soluble in 0.4 mol. LL¹ sodium chloride solution), prolamin (soluble in 70% (v/v) ethanol), and glutelin (soluble in 0.1 mol. LL¹ sodium hydroxide) were measured after semi-exhaustive extractions of whole-grain flour from the native grains, and barley (control) proteins (mg/g dry base (db)). These same species were also tested for total protein content, expressed as mg/g db. Error bars represent standard deviations (n = 3).

In comparison, teff (*Eragrostis tef*) has a reported total protein content of 8.5–9.4 g/100 g (Shumoy et al., 2018), which is markedly lower than that of Curly Mitchell Grass and slightly lower than the mean values observed for the other native species in this study. Browntop millet (*Urochloa ramosa*) contains 13.37 g/100 g protein, placing it within the same range as Button Grass, Native Millet, and Weeping Grass, but still considerably below the high protein content observed in Curly Mitchell Grass (29.1 g/100 g).

#### 3.1.4. Total lipid

The lipid content of the native grains exhibited considerable variation between species (Table 1). The values obtained encompass the total lipid content reported by Williams et al. (2024) for an unspecified native grain on a wet weight basis (5.0 ± 0.1 g/100 g wb). Native Millet had the highest lipid content with a value of 8.0 ± 0.6 g/100 g db (*p* ≤ 0.001), suggesting its potential as a valuable source of dietary lipids. The variation in lipid content among the native grains highlights the potential for selecting specific grains based on nutritional requirements and dietary goals. For example, Weeping Grass had the lowest lipid content (2.0 ± 0.4 g/100 g db) and might hold value in specific dietary applications where lower lipid intake is desired.

The higher lipid content in Native Millet might be due to an apparently large embryo (Abedi et al., 2023). Further research into the specific lipid profiles and fatty acid compositions of these grains would provide deeper insights into their nutritional benefits and potential health impacts. Teff (*Eragrostis tef*) as a reported lipid content of 2.38 g/100 g (Zhu, 2018), closely matching the values of Button Grass and Weeping Grass, but substantially lower than the lipid-rich Native Millet.

#### 3.1.5. Ash

Ash content of the native and control grains varied 4-fold between species (Table 1). As ash content is an indicator of the total mineral content, the high ash content in Button Grass may suggest that the grain could be richer in essential minerals such as calcium, magnesium, potassium, and phosphorus. Browntop millet (*Urochloa ramosa*) contains 5.48 g/100 g of ash (Ponnapalli et al., 2023), which falls between the higher ash content observed in Button Grass (8.7 g/100 g) and the lower values reported for Native Millet (3.4 g/100 g) and Weeping Grass (2.5 g/100 g), suggesting a moderate total mineral content.

### 3.2. Protein analysis

#### 3.2.1. Osborne protein fractionation

Our data uncovered notable variation in the proportions of the Osborne protein fractions in the native grain species, with barley as a control (Fig. 1). Among the studied grains, the glutelins were consistently the predominant fraction, except in Native Millet, for which the albumins were dominant. Weeping Grass had the highest abundance of glutelin (51.8% of the total, including albumins, globulins, prolamins, and glutelins). This contrasts with teff (*Eragrostis tef*), where glutelin is the least abundant protein fraction, comprising only 0.3– 0.6% of the total protein, while globulins dominate the profile, making up 9.6–13% (Shumoy et al., 2018).

In a study on rice starch systems, the addition of glutelin was found to raise the pasting temperature, indicating that gelatinisation occurred at a higher temperature, while reducing the viscosity of the starch paste. Increasing glutelin concentrations also led to higher gel hardness and adhesiveness. These findings suggest that glutelin dominance may influence food processing by producing firmer, more cohesive textures and altering water-binding behaviour. Such properties could affect dough structure, texture, and stability in products like noodles, gluten-free baked goods, or starch-based gels (Baxter et al., 2014).

The abundance of globulin was the lowest among the Osborne protein fractions across the native and control grain species, with Button Grass consistently showing the lowest globulin proportion (2.2% of the total) (*p* ≤ 0.001) compared to the other studied grains. The only exception was Weeping Grass, for which globulins represented 17.4% and the prolamin fraction represented the lowest proportion of total protein (7.1%).

In our microscopy study of the same four Australian native grains, we used 2,3,5-triphenyltetrazolium chloride solution to highlight the embryo in longitudinally cut grains (orange to red staining) and acid fuchsin to highlight protein in longitudinal sections (red staining) (Abedi et al., 2023). In that research, Curly Mitchell Grass and Native Millet were estimated to have a notably larger embryo as a proportion of the total grain compared with the other species (including sorghum and wheat), with concentrated protein observed in the aleurone and sub-aleurone layers of the longitudinal sections. In our present investigation, we found that the albumin fraction, which is known to be mainly distributed in the embryo and aleurone layers of cereal grains (Koehler & Wieser, 2013), constituted the highest proportion of total grain protein in Native Millet and the second-highest in Curly Mitchell Grass (Fig. 1).

In wheat, barley, and rye, a higher ratio of prolamin to glutelin in the dough generally indicates a weaker protein structure with less elasticity and increased susceptibility to deformation of the loaf during the baking process (Girard & Awika, 2021). However, the relationship between the prolamin:glutelin ratio in flour and the dough characteristics of gluten-containing cereals such as wheat, barley, and rye may not be applicable to gluten-free cereals. Despite conducting a thorough literature review, we were unable to identify studies that investigated the relationship between the prolamin:glutelin ratio and dough properties of gluten-free cereals. Further research is essential to establish a clearer understanding of the dough rheological characteristics of the native grains in relation to the prolamin:glutelin ratio.

To support future investigation into the relationship between the prolamin:glutelin ratio and dough properties in native grains, standard methods such as the farinograph and Rapid Visco Analyser (RVA) are recommended. The farinograph provides data on dough development time, water absorption, and stability, while the RVA measures pasting temperature, peak viscosity, and breakdown (among other pasting properties), reflecting starch-protein interactions. The RVA requires only small sample amounts (∼3 g), making it suitable for limited grain material, whereas the farinograph typically needs around 300 g of flour. These techniques can offer valuable insights into the functional behaviour of native grain flours in baking applications (Saleh et al., 2016).

#### 3.2.2. Protein profiles of Osborne fractions

In all the native species, both high-and low-molecular-weight proteins were present in the albumin and globulin fractions. Conversely, the prolamin and glutelin fractions were characterised solely by low-molecular-weight proteins.

The albumin fraction of Button Grass and Curly Mitchell Grass exhibited a wide range of proteins, spanning from 10 to 150 kDa (Fig. 2). The Native Millet albumin fraction displayed major protein bands ranging from 10 to 75 kDa, comparable to the barley protein profile. In the prolamin fraction, Button Grass, Native Millet, and Curly Mitchell Grass shared a similar band pattern, predominantly around 20 kDa. For the glutelin fraction, Button Grass, Native Millet, and Curly Mitchell Grass exhibited proteins ranging from 10 to 20 kDa, whereas Weeping Grass showed major protein bands between 10 and 15 kDa.

**Figure 2.**
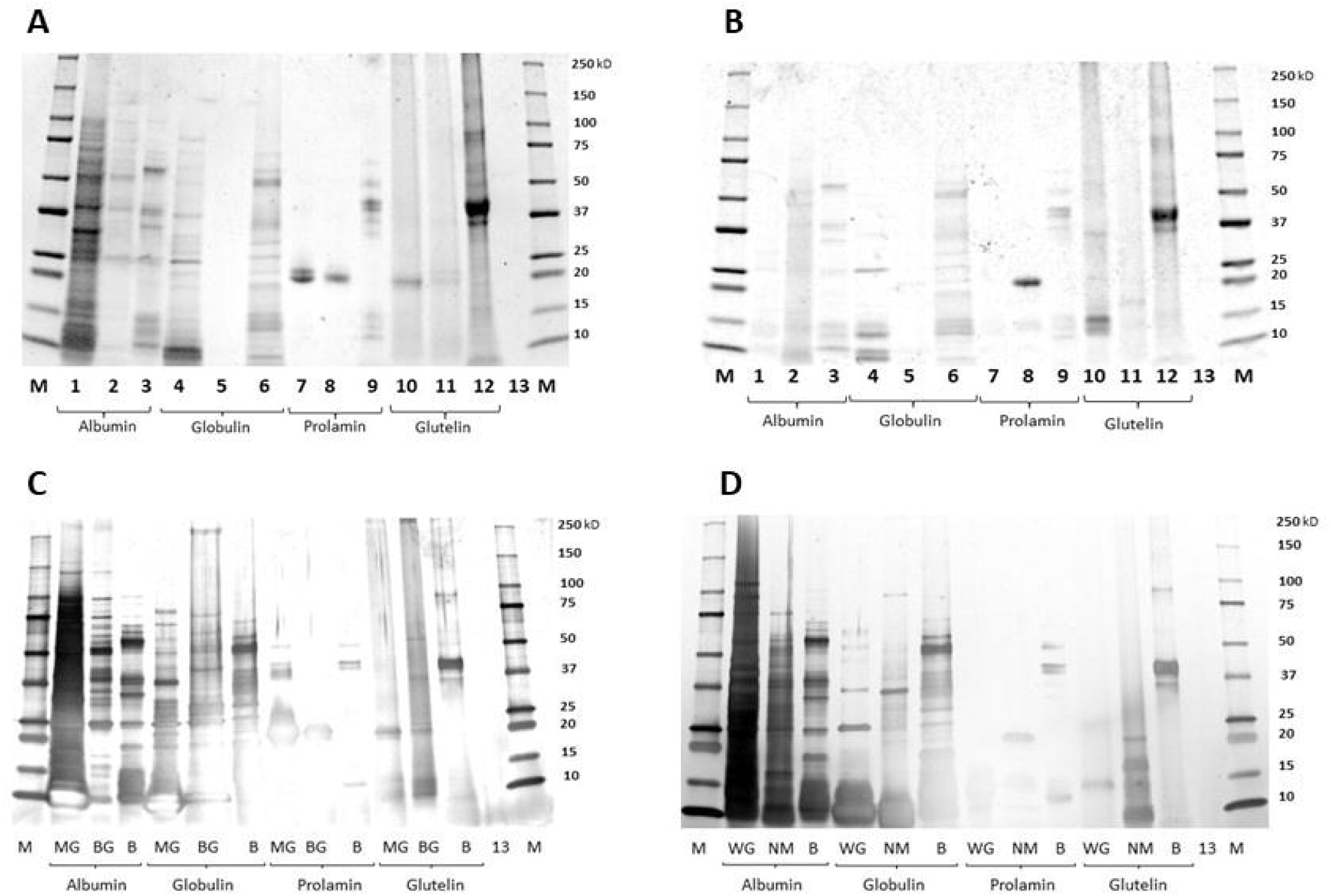
Protein profiles of the Osborne fractions in the native grains and barley. SDS-PAGE analysis showing the polypeptide profiles of albumin, globulin, prolamin, and glutelin fractions in Button Grass (BG), Curly Mitchell Grass (MG), Native Millet (NM), Weeping Grass (WG), and barley (B). The gels were stained with Coomassie Brilliant Blue (panels A, and B), and silver nitrate (panels C and D). Lanes M and 13 represent Marker, and blank, respectively.

SDS-PAGE analysis reported by Oom et al. (2008) confirmed the presence of α- and γ-kafirins in sorghum kafirin. The authors noted that γ-kafirin, which has a higher cysteine content than α-kafirin, is more likely to form disulfide bonds, contributing to progressive dough stiffening. While both these kafarins are low-molecular-weight proteins, their ability to cross-link can affect dough properties, particularly firmness. In contrast, wheat doughs owe their strength and elasticity to high-molecular-weight glutenin subunits, which form extensive polymeric networks (Shewry & Halford, 2002). This highlights a key functional difference in food applications: low-molecular-weight proteins may increase rigidity, but they do not support the extensibility or viscoelasticity provided by high-molecular-weight glutenins.

#### 3.2.3. Gluten screening

Our results confirmed that all the studied native grains—Button Grass, Curly Mitchell Grass, Native Millet, and Weeping Grass—were gluten-free, as indicated by the absence of a test line on the LFD (Fig.3). Among the control grains, only wheat and barley tested positive for gluten, which is consistent with their known gluten content. The observed faintness and incompleteness of the control band on the LFD for the barley sample can be attributed to the viscosity of the sample. The undiluted barley extract impacted the mobility of the buffer within the device, impeding its ability to ascend effectively to the top of the LFD. Conversely, sorghum and brown rice, also part of the control group, tested negative, consistent with their established gluten-free status (Fig. 3). The pink background observed on the LFD for the sorghum samples is caused by a natural red pigment present in the grain extract. Together, our results showing the gluten-free status of the native grains underscore the potential of using these species as safe alternatives in gluten-free diets.

**Figure 3.**
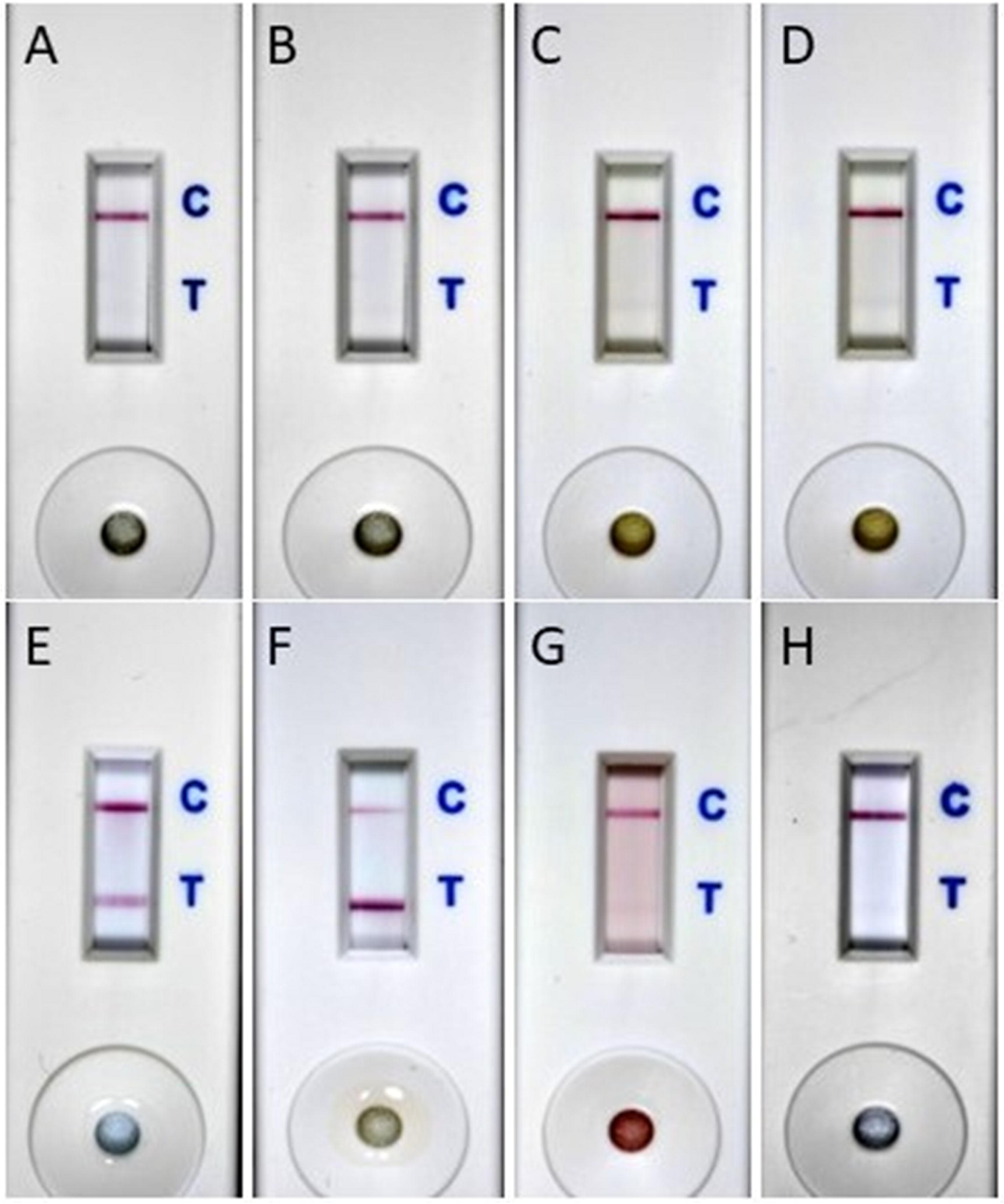
Gluten screening of the native and domesticated grains. The presence or absence of gluten in native and control grains was detected by a lateral flow device (section 2.4.3). A sample was considered positive for gluten if both the test and control lines were visible on the LFD; the absence of the test line or only a faint line indicated a negative result. The images are presented across eight panels: A (Button Grass), B (Curly Mitchell Grass), C (Native Millet), D (Weeping Grass), E (wheat), F (barley), G (sorghum), and H (brown Rice).

### 3.3. Carbohydrate profiles

#### 3.3.1. Total starch content

Total starch content among the studied grains was substantially lower on average compared to their domesticated counterparts (Table 2). The results were comparable to the pattern reported by Brand-Miller et al. (1998), in which native grains were found to contain 45.0 ± 2.0 g/100 g carbohydrate (combined starch and sugar). Our native grain total starch values, which ranged from 45.8 ± 0.4 g/100 g wb (equivalent to 53.7 ± 0.4 g/100 g db) for Native Millet to 53.5 ± 0.3 g/100 g wb (equivalent to 61.8 ± 0.3 g/100 g db) for Weeping Grass, were considerably lower than that reported for Kangaroo Grass (62.1 ± 0.8 g/100 g wb; Cowley et al., 2023). The total starch content of browntop millet (*Urochloa ramosum*) (56.43 ± 0.49 g/100 g db) falls within the range observed for the native grains in this study, being slightly higher than that of Native Millet and Button Grass, and close to Curly Mitchell Grass, but lower than Weeping Grass (Ponnapalli et al., 2023).

**Table 2.**
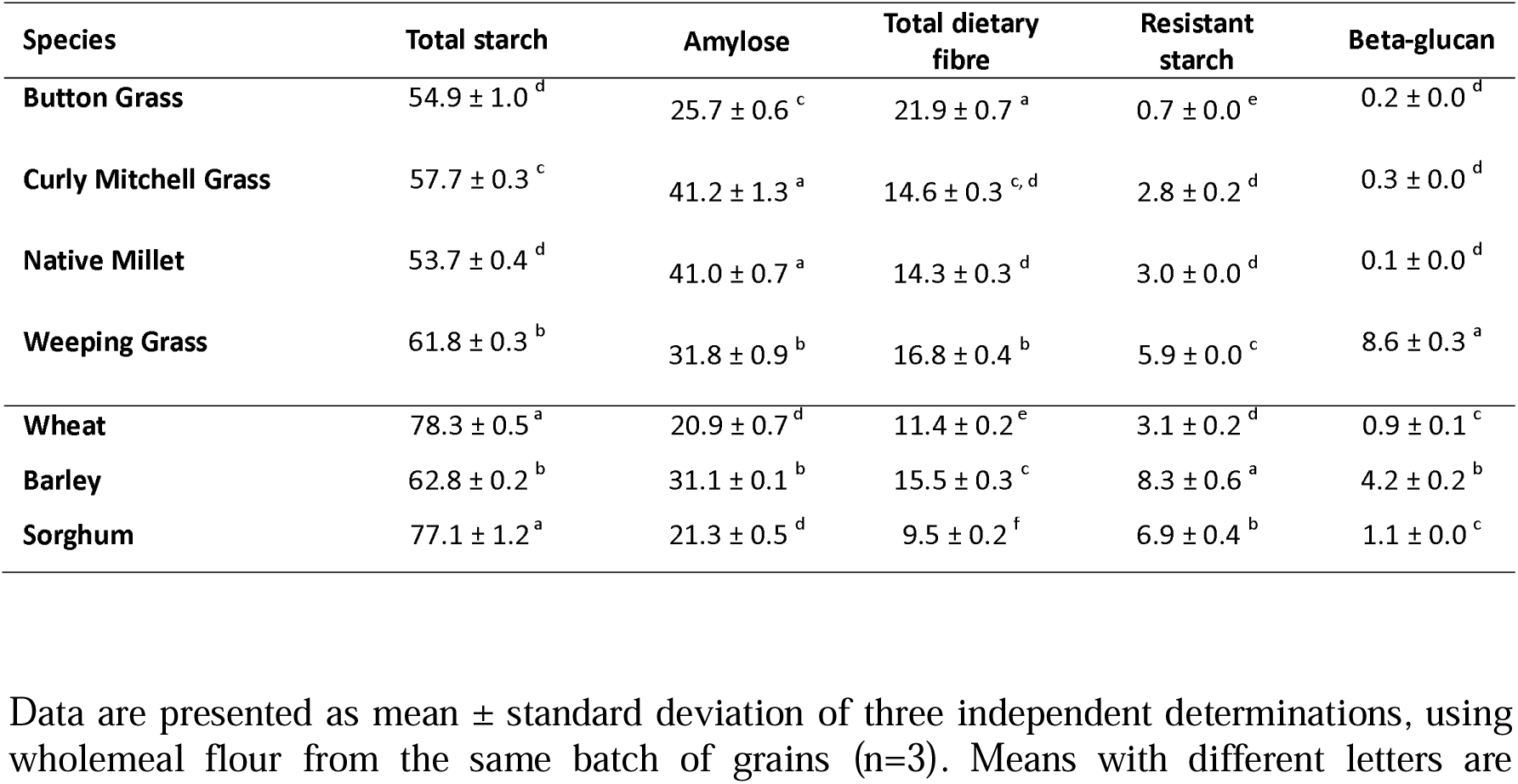
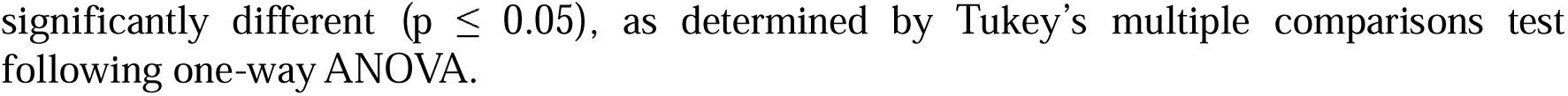
Total starch, amylose content, total dietary fibre, resistant starch, and beta-glucan of the native and control grains. All values are expressed as g/100 g (dry weight) of the grains except for amylose content, which is expressed as g/100 g of total starch.

| Species | Total starch | Amylose | Total dietary fibre | Resistant starch | Beta-glucan |
| --- | --- | --- | --- | --- | --- |
| Button Grass | 54.9 ± 1.0<sup>d</sup> | 25.7 ± 0.6<sup>c</sup> | 21.9 ± 0.7<sup>a</sup> | 0.7 ± 0.0<sup>e</sup> | 0.2 ± 0.0<sup>d</sup> |
| Curly Mitchell Grass | 57.7 ± 0.3<sup>c</sup> | 41.2 ± 1.3<sup>a</sup> | 14.6 ± 0.3<sup>c, d</sup> | 2.8 ± 0.2<sup>d</sup> | 0.3 ± 0.0<sup>d</sup> |
| Native Millet | 53.7 ± 0.4<sup>d</sup> | 41.0 ± 0.7<sup>a</sup> | 14.3 ± 0.3<sup>d</sup> | 3.0 ± 0.0<sup>d</sup> | 0.1 ± 0.0<sup>d</sup> |
| Weeping Grass | 61.8 ± 0.3<sup>b</sup> | 31.8 ± 0.9<sup>b</sup> | 16.8 ± 0.4<sup>b</sup> | 5.9 ± 0.0<sup>c</sup> | 8.6 ± 0.3<sup>a</sup> |
| Wheat | 78.3 ± 0.5<sup>a</sup> | 20.9 ± 0.7<sup>d</sup> | 11.4 ± 0.2<sup>e</sup> | 3.1 ± 0.2<sup>d</sup> | 0.9 ± 0.1<sup>c</sup> |
| Barley | 62.8 ± 0.2<sup>b</sup> | 31.1 ± 0.1<sup>b</sup> | 15.5 ± 0.3<sup>c</sup> | 8.3 ± 0.6<sup>a</sup> | 4.2 ± 0.2<sup>b</sup> |
| Sorghum | 77.1 ± 1.2<sup>a</sup> | 21.3 ± 0.5<sup>d</sup> | 9.5 ± 0.2<sup>f</sup> | 6.9 ± 0.4<sup>b</sup> | 1.1 ± 0.0<sup>c</sup> |

Data are presented as mean ± standard deviation of three independent determinations, using wholemeal flour from the same batch of grains (n=3). Means with different letters are
significantly different ( $p \leq 0.05$ ), as determined by Tukey's multiple comparisons test following one-way ANOVA.

#### 3.3.2. Total starch composition

Analysing the amylose contents of the studied grains revealed compelling variations in their starch composition. Curly Mitchell Grass and Native Millet, with amylose contents of 41.2 ± 1.3 and 41.0 ± 0.7 g/100 g of the total starch, respectively (Table 2), had the highest amylose content among the native and control grains (*p* ≤0.05). The high amylose content observed in these species might contribute to a relatively low glycaemic index, as amylose is closely linked to slowly digestible starch in cooked grains (Cowley et al., 2023).

Weeping Grass, with an amylose content of 31.8 ± 0.9 g/100 g of the total starch, aligned more closely with the control species, particularly barley (31.1 ± 0.1 g/100 g). Button Grass had the lowest amylose content among the native grains (25.7 ± 0.6 g/100 g), but somewhat higher than that observed in wheat (20.9 ± 0.7 g/100 g) and sorghum (21.3 ± 0.5 g/100 g) (Table 2).

The extent to which the ratio of amylose to amylopectin in the native grains will influence bread-making and the production of other grain-based foods needs to be further investigated. Flour (in wheat, for example) containing starch with a higher amylose:amylopectin ratio is known to absorb less water than flour with a lower amylose:amylopectin ratio (because amylose is less soluble in water than amylopectin) (Ee et al., 2020). This will also be true of the native grains, but, because their other components (such as lipids and proteins) are different, the effects of different amylose contents on product quality will have to be determined empirically.

#### 3.3.3. Total dietary fibre

The total dietary fibre content of Weeping Grass (16.8 ± 0.4 g /100 g) was similar to that of barley (Table 2). Curly Mitchell Grass exhibited a total dietary fibre content of 13.1 ± 0.2 g/100 g wb (equivalent to 14.6 ± 0.3 g/100 g db), which is comparable to the results reported by Foster et al. (2010), who found a total dietary fibre content of 15.0 g/100 g wb for Curly Mitchell Grass. Button Grass demonstrated a notably higher total dietary fibre content (21.9 ± 0.7 g/100 g) compared to barley and the other native grains analysed (Table 2; *p* ≤ 0.05). Teff (*Eragrostis tef*) has been reported to contain 8.0 g/100 g of total dietary fibre (Zhu, 2018), which is lower than the values observed in the native grains examined in this study, particularly Button Grass (21.9 g/100 g) and Weeping Grass (16.8 g/100 g), but closer to Curly Mitchell Grass (14.6 g/100 g) and Native Millet (14.3 g/100 g).

A higher total dietary fibre content can enhance the potential for breadmaking by increasing water absorption and mixing tolerance while decreasing extensibility. The higher fibre content may also extend the shelf-life of the bread by retaining moisture and slowing staling (Bagheri & Seyedein, 2011). Moreover, consumption of grains rich in dietary fibre is associated with reduced risks of non-communicable diseases including cardiovascular diseases, cancers, gastrointestinal disorders, obesity, and type 2 diabetes (Hojsak et al., 2022).

#### 3.3.4. Resistant starch

The native grains in our study consistently displayed lower levels of resistant starch compared to the control grains, particularly wheat (Table 2). The low resistant starch content of Button Grass (0.6 ± 0.0 g/100 g wb, equivalent to 0.7 ± 0.0 g/100 g db) was consistent with that reported in a previous study for Kangaroo Grass (0.9% w/w wb; Cowley et al., 2023).

Among the native grains, Weeping Grass exhibited the highest level of resistant starch (5.9 ± 0.0 g/100 g), indicating its potential for incorporation in baked goods, pasta, and beverages to improve texture and provide health benefits. Breads with an optimum resistant starch content exhibit increased crumb moisture, reduced firmness, and a lower retrogradation rate. The latter, coupled with the higher crumb moisture and water-retention capability of resistant starch, contributes to an overall reduction in crumb firmness in these breads. Resistant starch is also known to prolong the freshness of bread (Arp et al., 2021).

Weeping Grass also exhibited potential for utilisation in thickened, opaque health beverages, due to the swelling capacity and ability to form gels of resistant starch. In contrast to insoluble fibre, resistant starch provides a smoother mouthfeel and has a milder impact on sensory properties such as flavour (Tabibloghmany & Ehsandoost, 2014).

#### 3.3.5. Beta-glucan

The beta-glucan contents of the native grains in this study were higher than those reported for Kangaroo Grass (0.05 ± 0.0%, w/w) (Cowley et al., 2023). Weeping Grass displayed a higher beta-glucan content (7.47 ± 0.28 g/100 g wb, equivalent to 8.6 ± 0.3 g/100 g db) compared to both the native and control grains (*p* ≤ 0.0001) (Table 2). This finding builds on our microscopy-based study in which we identified that Weeping Grass had thicker endosperm cell walls than the other species investigated (Abedi et al., 2023), as evidenced by blue staining with calcofluor white, indicative of beta-glucan content.

Beta-glucan is an important dietary component, playing a key role in preventing cardiovascular diseases, managing obesity and diabetes, and regulating cholesterol levels in the body (Eraniappan et al., 2023). Using beta-glucan for its functional properties in foods, including thickening, stabilising, emulsifying, and gelling, offers opportunities to enhance ingredient stability, particularly in the production of beverages. When beta-glucans are incorporated into various products such as bread, muffins, pasta, noodles, salad dressings, beverages, soups, and reduced-fat dairy and meat items, it becomes apparent that their attributes, including breadmaking performance, water binding, emulsion stabilising capacity, thickening ability, texture, and appearance, are linked to the concentration, molecular weight, and structure of this polysaccharide (Kalinga & Mishra, 2009; Kaur et al., 2019; Sharma & Gujral, 2014).

### 3.4 Implications of the properties of the native grains for their potential in modern foods

The following highlights were observed regarding grain properties that distinguish these species from each other and/or their domesticated counterparts. The high lipid content of Native Millet may support its use in high-energy snack products. The high protein content and low carbohydrate content of Curly Mitchell Grass renders this grain potentially suitable for protein-enriched gluten-free products. We confirmed that all four of the native grains are gluten-free, reinforcing their suitability for gluten-free products such as breads and crackers. The large variation in amylose content among the native grains, including high levels in Curly Mitchell Grass and Native Millet, may enhance textural stability in products like noodles or baked goods, and support a lower glycaemic response due to the slower digestibility of high-amylose starches. The high dietary fibre content and low level of resistant starch in Button Grass aligns with its use in high-fibre cereal bars or biscuits. Finally, the high level of beta-glucan in Weeping Grass makes it a promising candidate for cholesterol-lowering breakfast cereals or functional beverages.

### 3.5 Environmental effects on grain quality

This study did not consider environmental variation, as all samples originated from specific locations and single seasons due to limited availability of these grains. Environmental factors—including rainfall, soil type, temperature, and duration of light exposure—can considerably influence grain composition (Endalamaw et al., 2025; Tester & Karkalas, 2001). Therefore, evaluating genotype × environment interactions is an important area for future research on Australian native grains, since it remains unclear whether these biochemical profiles are consistently expressed under different agronomic conditions.

## 4. Conclusion

This study has provided insights into the grain protein and carbohydrate complement of four species of Australian native grasses compared to their domesticated counterparts. These results have important implications for food applications of the native grains and human nutrition. Potential applications include partial substitutes for wheat in cereal-based foods (breads, pasta, noodles, snack bars) and beverages, as well as gluten-free baked products. However, the grain properties observed here cannot be extrapolated to all genotypes within these species or all plant growth environments. Understanding the relationships between grain properties and genotype × environment × management interactions for Australian native grains is the theme of our current research.

Consideration of the potential of native grains extends beyond the laboratory, with broad implications for the food industry, food production from hot and dry areas, and the empowerment of Indigenous communities. Incorporating these grains into food production not only has the potential to introduce valuable nutritional components and processing properties but also presents opportunities for sustainable economic activities, particularly for Indigenous communities, as well as revitalisation of cultural practices.

## CRediT authorship contribution statement

**Farkhondeh Abedi**: Formal analysis, Investigation, Methodology, Writing – original draft. **Claudia Keitel**: Supervision, Writing – review and editing. **Ali Khoddami**: Methodology, Supervision, Writing – review and editing. **Angela L. Pattison**: Resources, Writing – review and editing. **Thomas H. Roberts**: Conceptualization, Methodology, Project administration, Resources, Supervision, Writing – review and editing

## Acknowledgements

We thank Gamilaraay knowledge holders Kerrie Saunders, Hannah Binge, Diane Hall, and the other members of the Gamilaraay Grain Custodians, for sharing their knowledge of native grain cultivation and foods. We thank Jessica Maley, Iona Gyorgy, Hero Tahaei, Huajuan Ling, and Farhad Masoomi-Aladizgeh from the University of Sydney for assisting with the total protein, total dietary fibre, total lipid, and statistical analysis.

## Data availability

Data will be made available on request.

